# Efficient and multiplexed somatic genome editing with Cas12a mice

**DOI:** 10.1101/2024.03.07.583774

**Authors:** Jess D. Hebert, Haiqing Xu, Yuning J. Tang, Paloma A. Ruiz, Colin R. Detrick, Jing Wang, Nicholas W. Hughes, Oscar Donosa, Laura Andrejka, Saswati Karmakar, Irenosen Aboiralor, Rui Tang, Julien Sage, Le Cong, Dmitri A. Petrov, Monte M. Winslow

**Affiliations:** Department of Genetics, Stanford University School of Medicine, Stanford, CA, USA; Department of Radiology, Stanford University School of Medicine, Stanford, CA, USA; Department of Biology, Stanford University, Stanford, CA, USA; Cancer Biology Program, Stanford University School of Medicine, Stanford, CA, USA; Department of Pediatrics, Stanford University School of Medicine, Stanford, CA, USA; Department of Pathology, Stanford University School of Medicine, Stanford, CA, USA; Chan Zuckerberg Biohub Investigator

## Abstract

Somatic genome editing in mouse models has increased our understanding of the *in vivo* effects of genetic alterations in areas ranging from neuroscience to cancer biology and beyond. However, existing models are limited in their ability to create multiple targeted edits. Thus, our understanding of the complex genetic interactions that underlie development, homeostasis, and disease remains incomplete. Cas12a is an RNA-guided endonuclease with unique attributes that enable simple targeting of multiple genes with crRNA arrays containing tandem guides. To accelerate and expand the generation of complex genotypes in somatic cells, we generated transgenic mice with Cre-regulated and constitutive expression of enhanced *Acidaminococcus sp.* Cas12a (enAsCas12a). In these mice, enAsCas12a-mediated somatic genome editing robustly generated compound genotypes, as exemplified by the initiation of diverse cancer types driven by homozygous inactivation of trios of tumor suppressor genes. We further integrated these modular crRNA arrays with clonal barcoding to quantify the size and number of tumors with each array, as well as the efficiency of each crRNA. These Cas12a alleles will enable the rapid generation of disease models and broadly facilitate the high-throughput investigation of coincident genomic alterations in somatic cells *in vivo*.

## INTRODUCTION

Genetically engineered mouse models have been widely used to uncover phenotypes resulting from defined genetic alterations^1^. However, the creation of new mouse alleles has remained a barrier to the study of more complex genotypes, due to the expense and time required to generate a new allele and cross it with other alleles of interest^2^. Somatic genome editing with Cas9 has greatly increased the rate at which single genes can be studied both *ex vivo* and *in vivo*, but the ability of Cas9 to model complex genotypes has been limited by the need for each guide to have its own promoter and tracrRNA, thereby requiring the laborious cloning of guides in sequence^3^.

Cas12a (previously known as Cpf1) is a Class 2 Type V CRISPR-Cas system that has unique value in genome editing due to its ability to easily target multiple genomic loci^4–6^. Unlike Cas9, Cas12a has both RNase and DNase activity, and can process a single pre-crRNA transcript (crRNA array) containing numerous spacers (guides) into its constituent crRNAs to direct Cas12a-mediated cleavage of their target regions^4^. Cas12a is also distinct from Cas9 in recognizing a T-rich 4 bp protospacer adjacent motif (PAM) and generating staggered cuts that are distal to the PAM^5^. Cas12a from multiple species has been employed for genome editing *in vitro*, including Cas12a from *Acidaminococcus sp.* (As) and *Lachnospiraceae bacterium* (Lb)^4,6–11^. Large-scale screens have identified Cas12a variants with increased efficiency, broadened PAM recognition ranges, and reduced off-target cleavage^12–15^. In particular, an enhanced version of AsCas12a (enAsCas12a) with several substitutions (E174R/N282A/S542R/K548R) has superior genome editing relative to wild type AsCas12a, in addition to a substantially expanded targeting range^13^.

The impact of complex genotypes is highly relevant in human cancer, which is defined by its genetic complexity. Individual tumors can have tens to hundreds of non-synonymous mutations, in addition to aberrant epigenetic modifications, gene duplications, chromosomal rearrangements, and gains or losses of entire chromosomes^16–18^. Any or all of these alterations could contribute to tumorigenesis, tumor progression, and resistance to therapy. Deconvoluting functional drivers from human cancer sequencing data and understanding how these genes interact to generate cancer phenotypes is complicated by the limited availability of samples, differences among patients and which therapies they have received, and technical variability of the approaches used to analyze them. Functional interrogation of the phenotypes resulting from defined combinations of alterations is thus critical for isolating the key drivers of these complex disease states.

To accelerate and expand our ability to generate complex genotypes in somatic cells, we generated Cre-regulated and constitutive enAsCas12a transgenic mice. Efficiently multiplexed somatic genome editing using Cas12a transgenic mice enables the rapid generation of complex models and should facilitate the high-throughput investigation of coincident genomic alterations *in vivo*.

## RESULTS

### Generation and validation of Cre-regulated and constitutive enAsCas12a alleles

To enable Cas12a-mediated somatic genomic editing and thus take advantage of the features that distinguish Cas12a from Cas9 (**Supplementary Figure 1a**), we generated transgenic mice expressing enAsCas12a with an optimized nuclear localization sequence (NLS) configuration that increases cutting efficiency and an HA tag to facilitate identification (enAsCas12a; **Figure 1a**)^13,19^. We generated transgenic mice through integration of a CAGGS promoter-driven Lox-Stop-Lox (LSL)-enAsCas12a-PolyA cassette into the *H11* locus (*H11^LSL-Cas12a^*) (**Figure 1a,b**). Expression of Cre in fibroblasts from *H11^LSL-Cas12a^* mice induced recombination of the LSL cassette and expression of Cas12a protein (**Figure 1c**). We also crossed *H11^LSL-Cas12a^* mice to CMV-Cre “deleter” mice to generate a constitutive *H11^Cas12a^* allele (**Figure 1d** and **Supplementary Figure 1b**). *H11^Cas12a^* mice were viable and fertile and had widespread expression of Cas12a (**Figure 1e**), with prominently nuclear protein localization (**Figure 1f** and Supplementary Figure 1c).

**Figure 1.**
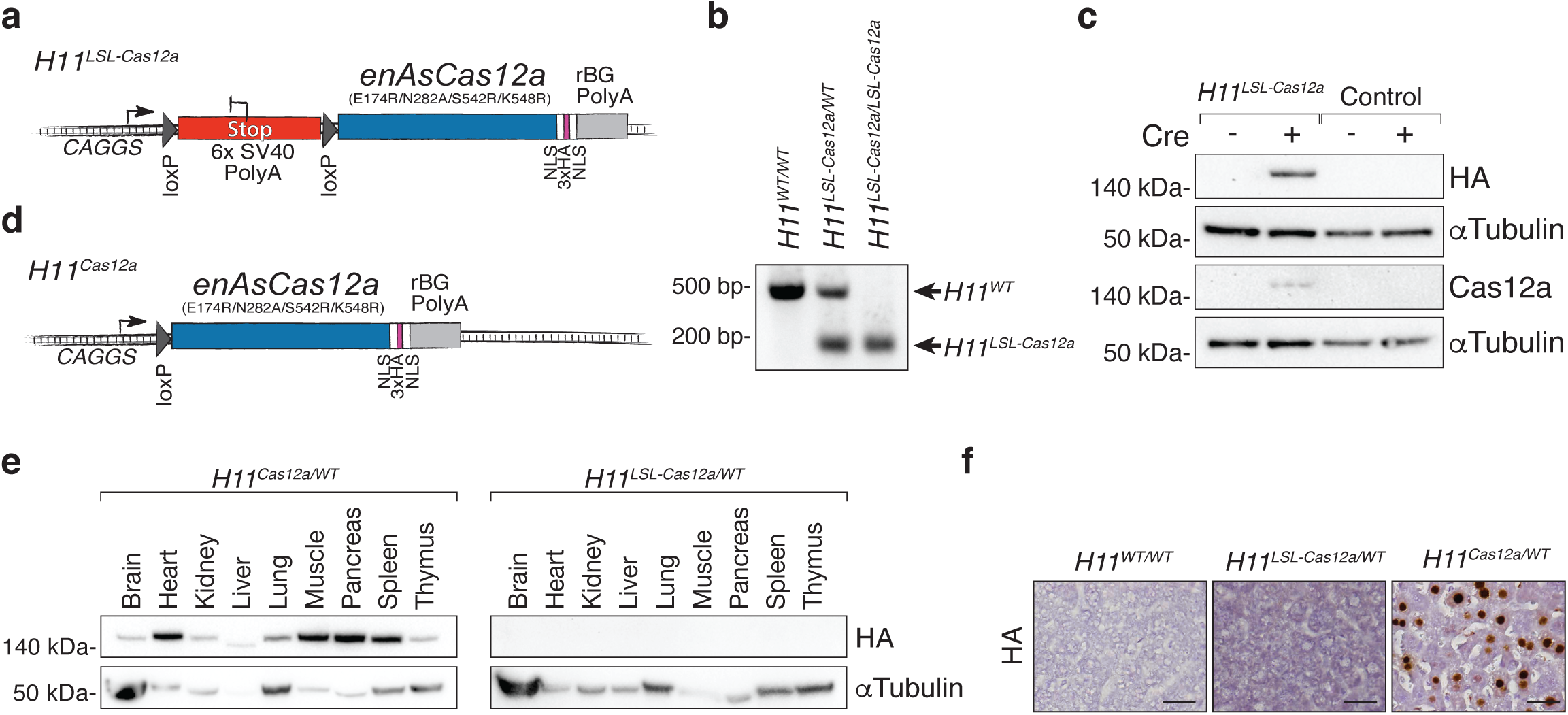
Generation of Cre-regulated and constitutive Cas12a mice. **a.** Schematic of the *H11^LSL-Cas12a^* transgene. Enhanced AsCas12a containing several substitutions to increase on-target efficiency and PAM binding sequence range (Kleinstiver *et al.* (2019). *Nature Biotechnology*), has two nuclear localization sequences (NLS), and three HA tags at the C-terminus (Liu *et al.* (2019), *Nucleic Acids Research*), controlled by Cre-mediated removal of the LoxP-Stop-LoxP (LSL) cassette. **b.** PCR genotyping of mice of the indicated *H11^LSL-Cas12a^* and wild-type (*WT*) genotypes. **c.** Western blots on tail tip fibroblasts from *H11^LSL-Cas12a^* and wild-type control mice 3 days after Adeno-Cre (Cre) treatment. Cas12a was detected by both anti-HA and anti-Cas12a antibodies. aTubulin shows loading. **d.** Schematic of the *H11^Cas12a^* transgene, following Cre-mediated removal of the LSL cassette. **e.** Western blots on tissue lysates from *H11^Cas12a/WT^* or *H11^LSL-Cas12a/WT^* mice. a-Tubulin shows loading. **f.** Immunohistochemical staining for the HA tag on Cas12a on liver sections from the indicated genotypes of mice. Scale bars, 25 μm.

### Cas12a-induced inactivation of *Nf1, Rasa1* and *Pten* generate lung adenocarcinoma

Oncogene-negative lung adenocarcinoma represents ∼30% of all lung adenocarcinomas and has been modeled in mice through combinatorial inactivation of the tumor suppressor genes *Nf1*, *Rasa1* and *Pten* in lung epithelial cells^20^. This was initially accomplished using Cas9-mediated somatic genome editing and multi-step sgRNA cloning^20^. As an initial test of the ability of the *H11^LSL-Cas12a^* allele to generate tumors with complex genotypes *in vivo*, we generated lentiviral vectors expressing Cre and a pre-crRNA array targeting *Nf1*, *Rasa1*, and *Pten*. Given the ability to synthesize these three-crRNA arrays and their ease of cloning, we generated a pool of 27 lentiviral Cre vectors with every combination of three guides targeting each tumor suppressor gene (**Figure 2a**). We cloned these pre-crRNA arrays into a vector that contained a diverse 16-nucleotide barcode (BC) at the 3’ end of the U6 promoter and Cre recombinase (Lenti-U6^BC^-cr*Nf1/Rasa1/Pten*-Cre; **Figure 2b**). Cre in these vectors induces Cas12a expression in somatic cells in *H11^LSL-Cas12a^* mice, and amplification of the BC-crRNA region from bulk tumor-bearing lungs followed by tumor barcoding and high-throughput barcode sequencing (Tuba-seq^Ultra^; U6 barcode Labeling with per-Tumor Resolution Analysis) can quantify the size of each clonal tumor^21–23^.

**Figure 2.**
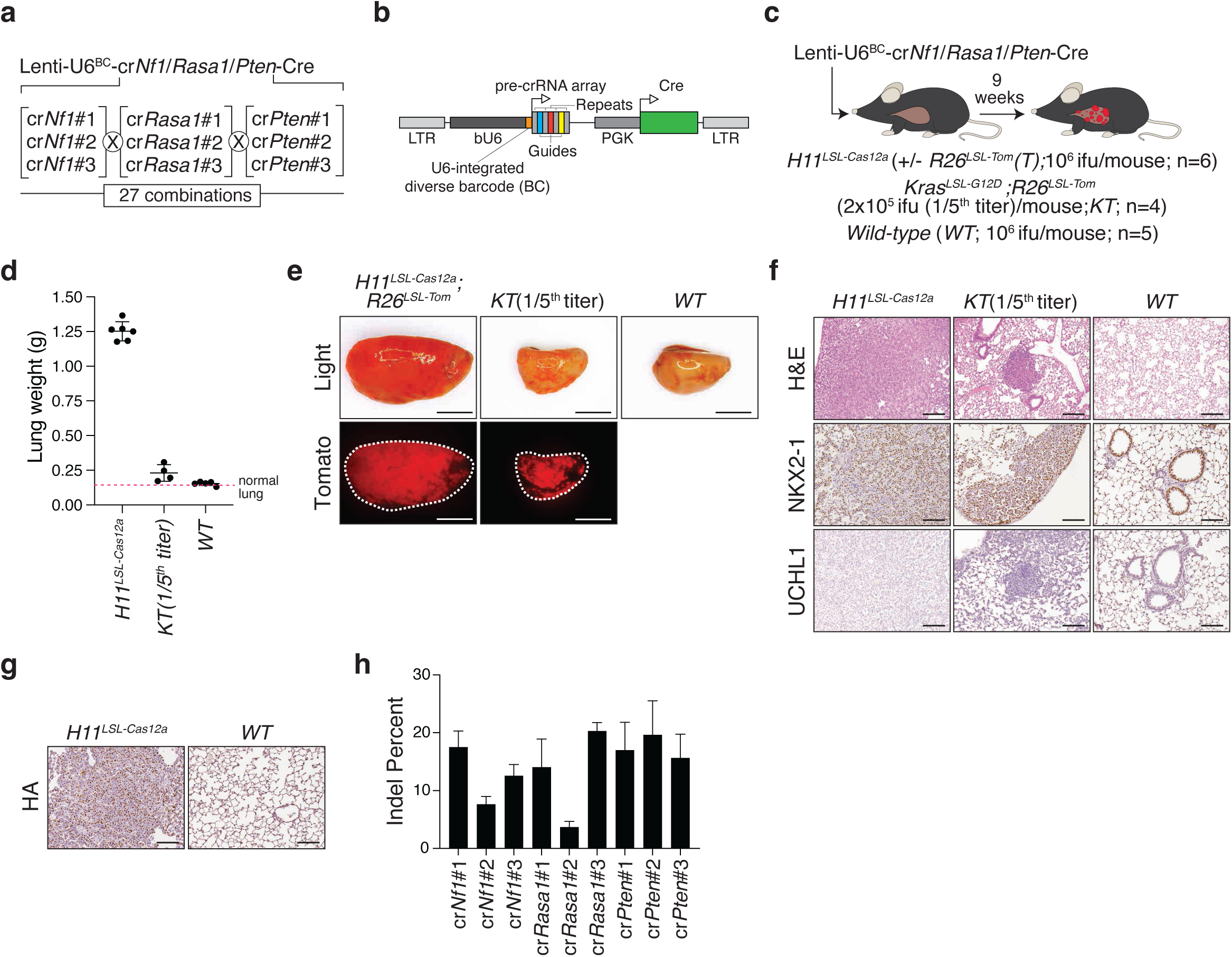
Rapid and efficient generation of oncogene-negative lung tumors through Cas12a-mediated coincident inactivation of three tumor suppressor genes. a. Lenti-U6^BC^-cr*Nf1/Rasa1/Pten*-Cre pool has guides targeting *Nf1, Rasa1*, and *Pten*. Each gene is targeted by three crRNAs in all combinations. b. Design of a lentiviral vector that expresses Cre recombinase and has a pre-crRNA array with three spacers (guides) downstream of a bovine U6 promoter with an integrated barcode region. c. Intratracheal delivery of Lenti-U6^BC^-cr*Nf1/Rasa1/Pten*-Cre to *H11^LSL-Cas12a^, Kras^LSL-G12D^;R26^LSL-Tom^* (*KT*), and *wild-type (WT)* mice. Viral titer (infectious units, ifu) and mouse numbers are indicated. d. Lung weights from mice of the indicated genotypes 9 weeks after transduction with Lenti-U6^BC^cr*Nf1/Rasa1/Pten*-Cre. Each dot represents a mouse. Means +/- standard deviation are indicated. e. Light (upper) and Tomato fluorescent (lower) images of lung lobes from the indicated genotypes of mice 9 weeks after transduction with Lenti-U6^BC^-cr*Nf1/Rasa1/Pten*-Cre. Dashed lines outline tissues. Tumors are Tomato-positive due to recombination of the *R26^LSL-Tomato^* allele. Scale bars, 5 mm. f. Hematoxylin & eosin (upper), NKX2-1 (middle), and UCHL1 (lower) staining of lung sections from the indicated genotypes. Scale bars, 100 μm. g. Immunohistochemical staining for the Cas12a HA tag on lung tissue from representative *H11^LSL-Cas12a^* and *WT* mice transduced with Lenti-U6^BC^-cr*Nf1/Rasa1/Pten*-Cre. Scale bars, 100 μm. h. Indel frequencies for genomic regions targeted by each crRNA within bulk tumor-bearing lung tissue from *H11^LSL-Cas12a^* mice. Means +/- standard deviation of three tumor-bearing lungs are shown.

We transduced the lungs of *H11^LSL-Cas12a^* mice (some of which also contained a *R26^LSL-Tomato^* Cre-reporter allele), wild-type negative control mice, and *Kras^LSL-G12D^;R26^LSL-Tomato^*(*KT*) positive control mice with Lenti-U6^BC^-cr*Nf1/Rasa1/Pten*-Cre (**Figure 2c**). Tumors in *KT* mice form due to Cre-mediated expression of oncogenic KRAS in the absence of any Cas12a-mediated gene targeting. We anticipated that Cas12a-mediated homozygous inactivation of *Nf1, Rasa1,* and *Pten* might be limited; therefore, we used a 5-fold higher titer in *H11^LSL-Cas12a^* mice relative to *KT* mice in which every transduced cell will express oncogenic KRAS. Unexpectedly, only 9 weeks after tumor initiation, the *H11^LSL-Cas12a^* mice developed extremely high tumor burden as assessed by lung weight (>15-folder higher than *KT* mice after subtracting normal lung weight), direct imaging of the lung lobes, and histology (**Figure 2 d-f**). Histology, positive immunohistochemical staining for NKX2-1/TTF-1, and negative staining for UCHL1 confirmed that these tumors were lung adenomas and adenocarcinomas (**Figure 2f**)^20^. Neoplastic cells in these tumors expressed nuclear Cas12a (**Supplementary Figure 2g**). PCR amplification and Sanger sequencing of each of the 9 targeted regions from genomic DNA from bulk tumor-bearing lungs from *H11^LSL-Cas12a^*mice confirmed indels at each target site (**Figure 2h**).

### Quantification of Cas12a-mediated tumorigenesis

We PCR-amplified and high-throughput sequenced the BC-crRNA region of the integrated Lenti-U6^BC^-cr*Nf1/Rasa1/Pten*-Cre vectors from bulk tumor-bearing lungs (Tuba-seq^Ultra^)^21,23^. This approach enabled the precise quantification of the number of cancer cells in each clonal tumor, the number of tumors in each mouse, and the relative efficiency of each guide (**Figure 3a**). Overall tumor burden normalized to viral titer was more than 10-fold greater in *H11^LSL-Cas12a^* than *KT* mice (**Figure 3b**). We also assessed the number of clonal barcoded tumors in each mouse and found that *H11^LSL-Cas12a^* mice had 2-fold more tumors on average than *Kras^LSL-G12D^* mice when corrected for viral titer (**Figure 3c**). These results are consistent with the high tumorigenic potential of lung epithelial cells with combined inactivation of *Nf1*, *Rasa1*, and *Pten*^20,24^.

**Figure 3.**
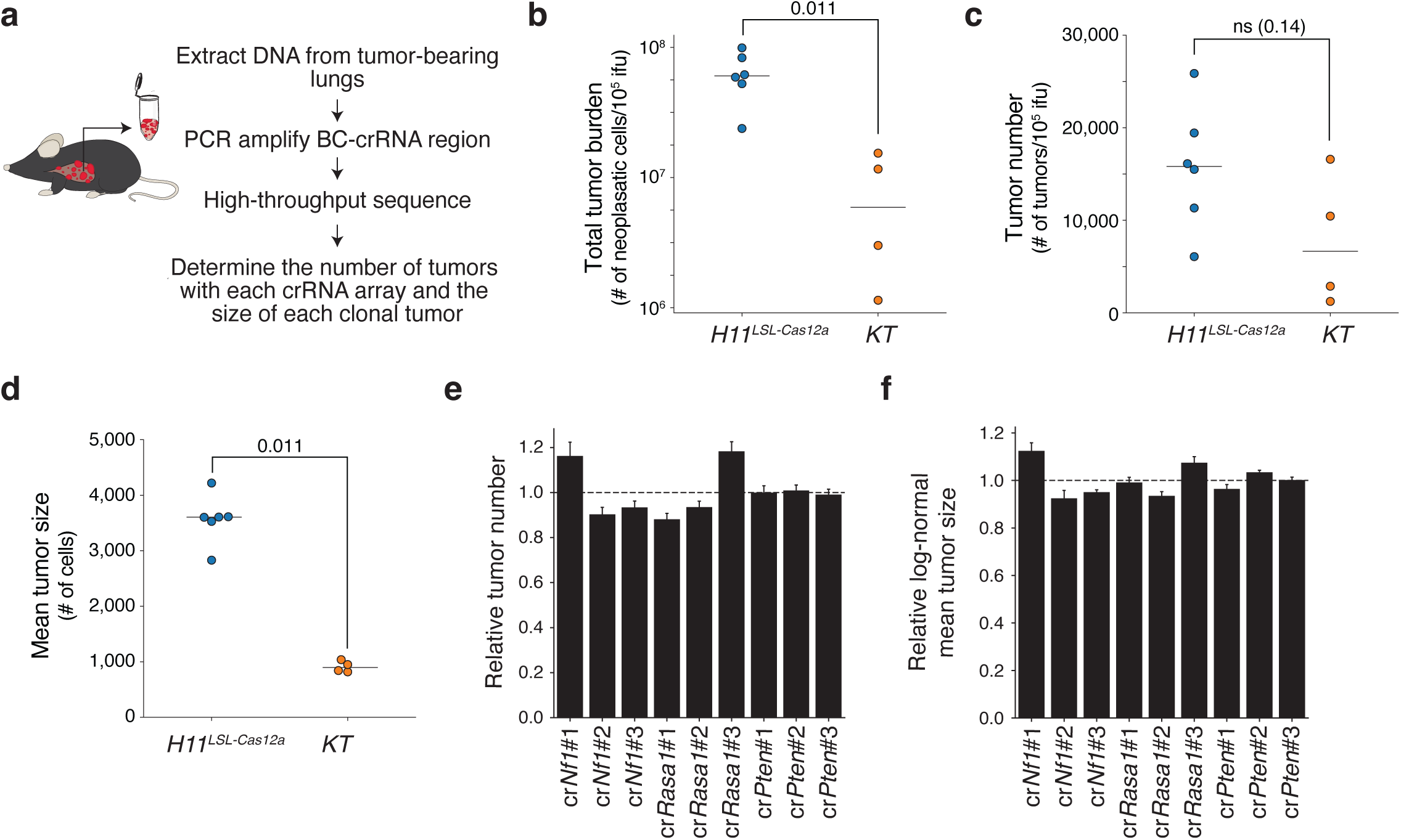
Integration of crRNA arrays with tumor barcoding enables quantification of tumor initiation and tumor size. a. Schematic of tumor barcoding with high-throughput BC-crRNA sequencing to determine the size of each Cas12a-induced clonal tumor. b. Total number of neoplastic cells (Total tumor burden) in each mouse normalized to viral titer. Each dot represents a mouse, and bars are means. P-value calculated with Wilcoxon rank-sum test. c. Number of tumors in each mouse normalized to viral titer. Each dot represents a mouse, and bars are means. P-value calculated with Wilcoxon rank-sum test. d. Mean tumor size given a log-normal tumor size distribution. Each dot represents a mouse, and bars are means. P-value calculat-ed with Wilcoxon rank-sum test. e. Number of tumors for each crRNA (guide) relative to the median value for each gene. 95% confidence intervals are shown. f. Mean tumor size assuming a log-normal distribution for each crRNA (guide) relative to the median value for each gene. 95% confidence intervals are shown.

Tumor sizes in *H11^LSL-Cas12a^* mice were dramatically larger than in *KT* mice, with the log-normal mean tumor size around 4-fold greater in *H11^LSL-Cas12a^* mice (**Figure 3d**), suggesting increased tumor growth of cr*Nf1/Rasa1/Pten* tumors compared to those driven by oncogenic KRAS. Finally, each of the 9 crRNAs (**Figure 3e-f**) and 27 crRNA arrays (**Supplementary Figure 2a-b**) generated relatively similar tumor numbers and sizes. These results indicate efficient gene inactivation and multiplexed somatic genome editing in *H11^LSL-Cas12a^* mice.

### Induction of small-cell lung cancer through the Cas12a-mediated inactivation of *Rb1, Trp53*, and *Rbl2*

Small-cell lung cancer (SCLC) is a neuroendocrine cancer that has been modeled in mice through Cre/lox-mediated, and more recently Cas9-mediated, inactivation of *Rb1*, *Trp53* and *Rbl2*^25,26^. We generated a lentiviral vector expressing Cre and a pre-crRNA array with all combinations of three guides targeting *Rb1*, *Trp53*, and *Rbl2* to create 27 vectors for combinatorial gene inactivation (Lenti-U6^BC^-cr*Rb1/Trp53/Rbl2*-Cre; **Figure 4a**). We transduced *H11^LSL-Cas12a^* mice (some of which also contained an *R26^LSL-Tomato^* allele) and wild-type negative control mice with Lenti-U6^BC^-cr*Rb1/Trp53/Rbl2*-Cre (**Figure 4b**). After 18 weeks, transduced *H11^LSL-Cas12a^* mice had high lung tumor burden (**Figure 4c-d**). Tumors in these mice included UCHL1^positive^ bronchiolar early-stage SCLC, along with more poorly differentiated regions (**Figure 4e**). Cancer cells stained positive for the Cas12a HA tag (**Figure 4e**). Amplification of the BC-crRNA region from bulk tumor-bearing lungs, high-throughput sequencing, and Tuba-seq^Ultra^ analysis revealed that *H11^LSL-Cas12a^* mice transduced with Lenti-U6^BC^-cr*Rb1/Trp53/Rbl2*-Cre had large numbers of clonal tumors and high tumor burden, consistent with robust tumor growth (**Figure 4f-g**). Microdissected tumors had indels at the targeted genomic sites (**Figure 4h**). Quantitative analysis of the crRNA arrays in tumors showed that different crRNAs generated similar tumor numbers and sizes (**Figure 4i-j**). Thus, *H11^LSL-Cas12a^* mice enabled rapid and efficient generation of this recalcitrant cancer type.

**Figure 4.**
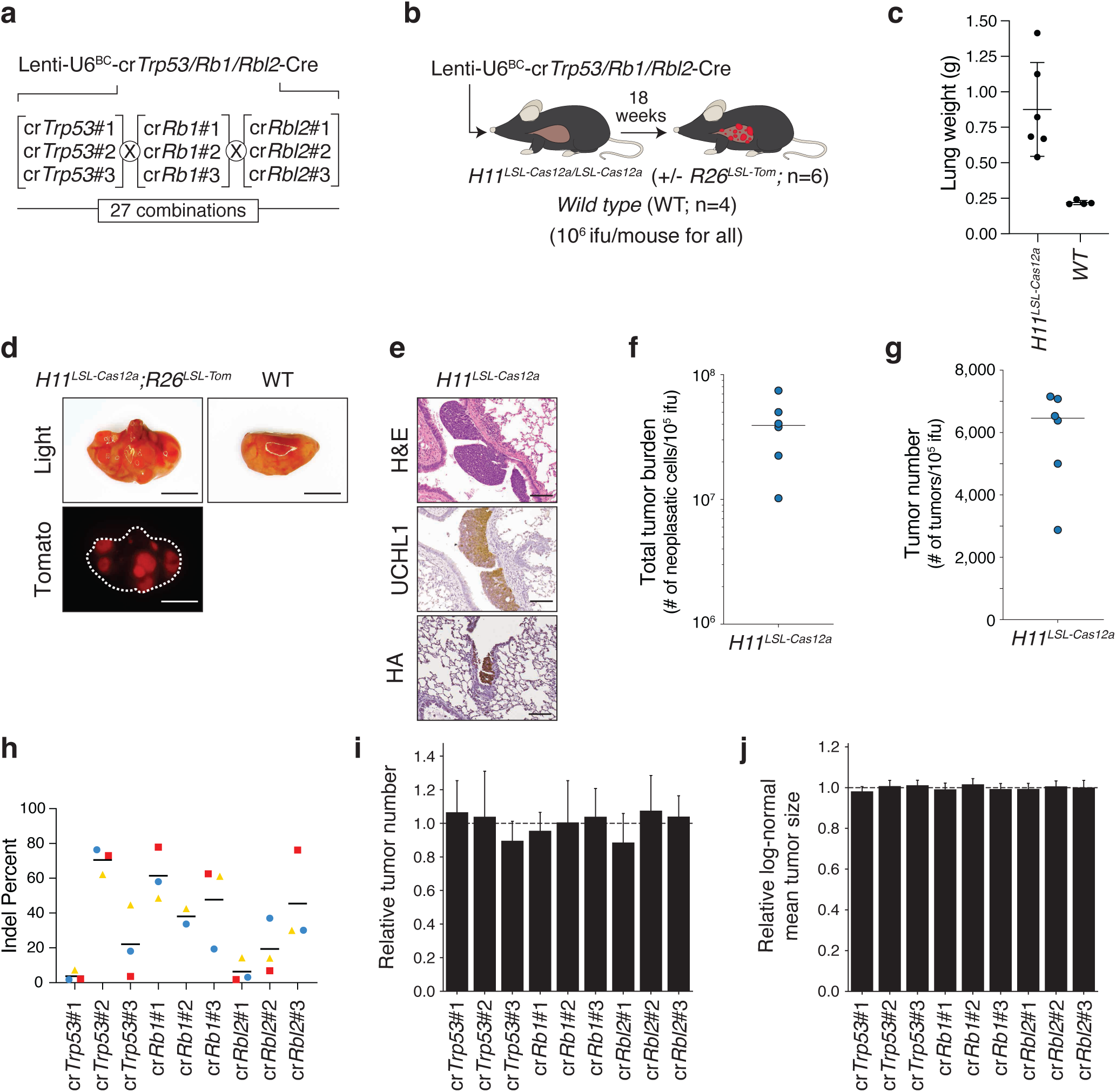
Induction of small-cell lung cancer through the simultaneous Cas12a-mediated inactivation of *Trp53, Rb1* and *Rbl2*. **a.** Lenti-U6^BC^-cr*Trp53/Rb1/Rbl2*-Cre pool has guides targeting *Trp53, Rb1*, and *Rbl2*. Each gene is targeted by three crRNAs in all 27 combinations. b. Intratracheal delivery of Lenti-U6^BC^-cr*Trp53/Rb1/Rbl2*-Cre to *H11^LSL-Cas12a^* and *wild-type (WT)* mice. Viral titer (infectious units, ifu) and mouse numbers are indicated. c. Lung weights from mice of the indicated genotypes 18 weeks after transduction with Lenti-U6^BC^-cr*Trp53/Rb1/Rbl2*-Cre. Each dot represents a mouse. Mean +/- standard deviation is indicated. d. Light (upper) and Tomato fluorescent (lower) images of lung lobes from the indicated genotypes of mice 18 weeks after transduction with Lenti-U6^BC^-cr*Trp53/Rb1/Rbl2*-Cre. Dashed line outlines tissue. Tumors are Tomato-positive due to recombination of the *R26^LSL-Tomato^* allele. Scale bars, 5 mm. e. Hematoxylin & eosin (upper), UCHL1 (middle) and HA tag (lower) staining of lung sections from an *H11^LSL-Cas12a^* mouse. Scale bars, 100 μm. f. Total number of neoplastic cells (Total tumor burden) in each mouse normalized to viral titer. Each dot represents a mouse, and the bar is the mean. g. The number of tumors in each mouse normalized to viral titer. Each dot represents a mouse, and the bar is the mean. h. Indel frequencies for genomic regions targeted by each crRNA within micro-dissected tumor tissue from *H11^LSL-Cas12a^* mice. Symbol shapes and colors identify values from the same sample. Bars represent means. **i,j.** Number of tumors (**i**) and mean tumor size assuming a log-normal distribution (**j**) for each crRNA (guide) relative to the median value for each gene. 95% confidence intervals are shown.

### Generation of autochthonous PDAC with inactivation of *Trp53*, *Cdkn2a*, and *Smad4*

*TP53*, *CDKN2A*, and *SMAD4* are the three most frequently mutated tumor suppressor genes in human pancreatic ductal adenocarcinoma (PDAC), and many diagrams of the genomic progression of this cancer type show the acquisition of mutations/alterations in all three of these genes^27,28^. However, despite extensive work to investigate these tumor suppressor genes in pancreatic carcinogenesis *in vivo*^29–35^, a model in which all three tumor suppressor genes are inactivated has yet to be published. To determine whether Cas12a-mediated somatic genome editing would also be efficient in adult pancreatic epithelial cells, we generated a lentiviral vector expressing Cre and pre-crRNA arrays with three gRNAs targeting *Trp53*, *Cdkn2a*, and *Smad4* (Lenti-U6^BC^-cr*Trp53/Cdkn2a/Smad4*-Cre; **Figure 5a**).

**Figure 5.**
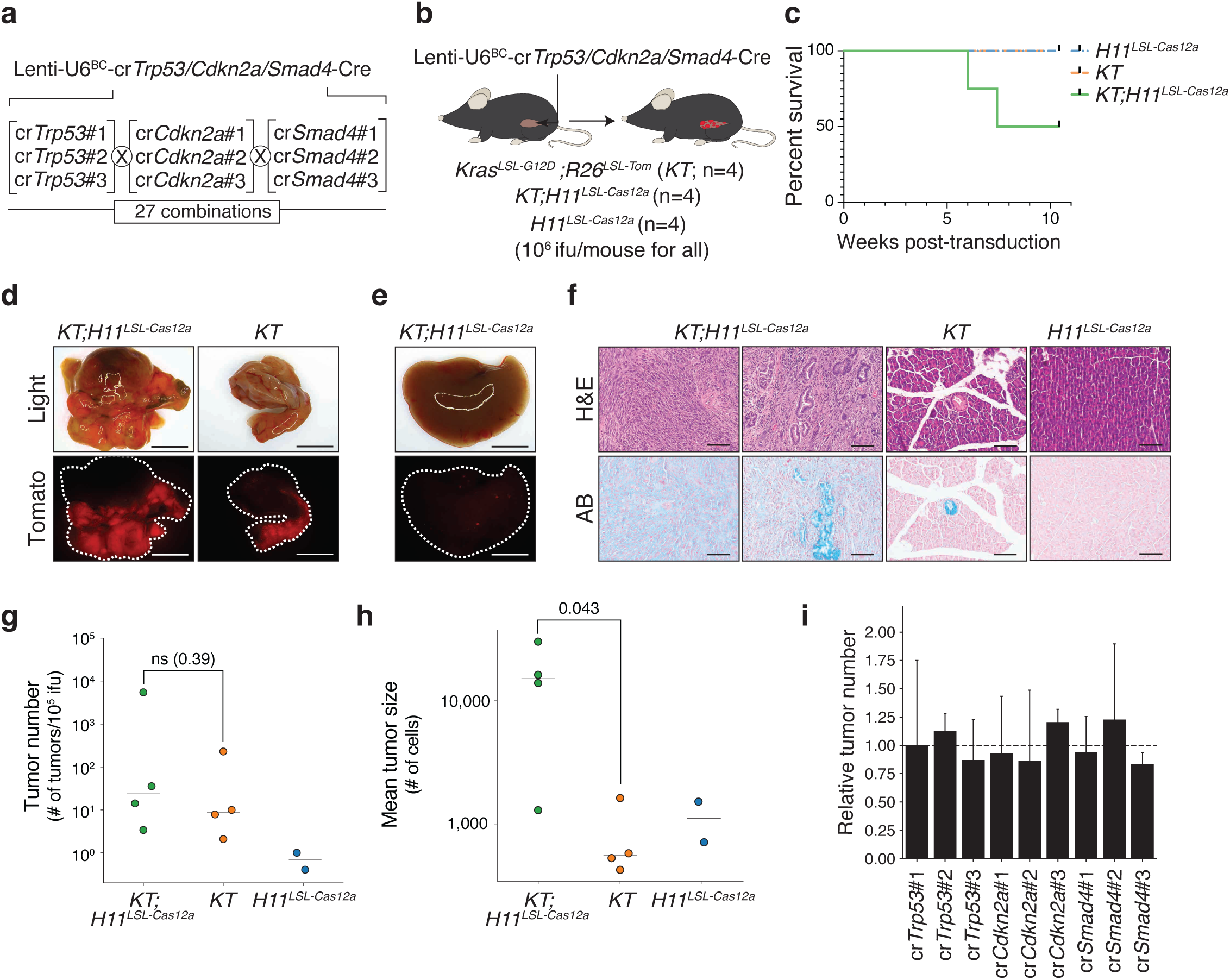
Rapid generation of PDAC through somatic Cas12a-mediated inactivation of commonly mutated tumor suppressor genes. a. Lenti-U6^BC^-cr*Trp53/Cdkn2a/Smad4*-Cre pool has guides targeting *Trp53, Cdkn2a*, and *Smad4*. Each gene is targeted by three crRNAs in all 27 combinations. b. Intrapancreatic delivery of Lenti-U6^BC^-cr*Trp53/Cdkn2a/Smad4*-Cre to *Kras^LSL-G12D^;R26^LSL-Tom^*(*KT*)*, KT;H11^LSL-Cas12a^*, and *H11^LSL-Cas12a^* mice. Viral titer (infectious units, ifu) and mouse numbers are indicated. c. Survival curve of mice of the indicated genotypes transduced with Lenti-U6^BC^-cr*Trp53/Cdkn2a/Smad4*-Cre. d. Light (upper) and fluorescent Tomato (lower) pancreas images from the indicated genotypes of mice 6 weeks (*KT;H11^LSL-Cas12a^*) or 11 weeks (*KT*) after transduction with Lenti-U6^BC^-cr*Trp53/Cdkn2a/Smad4*-Cre. Dashed lines outline tissues. Scale bars, 5 mm. e. Light and fluorescent Tomato images of a liver lobe from a *KT;H11^LSL-Cas12a^* mouse. Dashed line outlines tissue. Scale bars, 5 mm. f. Immunohistochemistry images of pancreas sections from the indicated genotypes, showing hematoxylin & eosin (H & E; upper) and Alcian Blue (AB; lower) staining. *KT;H11^LSL-Cas12a^* panels show representative regions with poorly differentiated sarcomatoid carcinoma (left; approximately 80-90% of tumor area) and more differentiated areas with mucinous cells and some lobular structure (right). Scale bars, 100 μm. g. The number of tumors in each mouse normalized to viral titer. Each dot represents a mouse, and bars are means. Note that two *H11^LSL-Cas12a^* mice did not have detectable tumor burden and are not plotted. P-value calculated with Wilcoxon rank-sum test. h. Mean tumor size given a log-normal tumor size distribution. Each dot represents a mouse, and bars are means. Note that two *H11^LSL-Cas12a^* mice did not have detectable tumor burden and are not plotted. P-value calculated with Wilcoxon rank-sum test. i. Number of tumors for each crRNA (guide) relative to the median value for each gene. 95% confidence intervals are shown.

We delivered this pool to the pancreata of *KT;H11^LSL-Cas12a^*, Cas12a-negative *KT*, and *H11^LSL-Cas12a^* mice through retrograde pancreatic ductal injection (**Figure 5b**)^36^. Several *KT;H11^LSL-Cas12a^* mice developed large multifocal pancreatic tumors as early as 6 weeks after transduction, with some also showing metastatic dissemination to the liver (**Figure 5c-e**). While *KT* mice only developed PanIN lesions, *KT;H11^LSL-Cas12a^* mice developed large, Cas12a-expressing primary tumors, including poorly-differentiated PDAC with cytological atypia and smaller Cytokeratin-19^positive^ mucinous regions with desmoplastic stroma and lobular structures (**Figure 5f** and **Supplementary Figure 3a-b**). The pancreata of transduced *H11^LSL-Cas12a^* mice appeared entirely normal, suggesting that even the combined inactivation of these three tumor suppressors has minimal effect in the absence of oncogenic KRAS (**Figure 4f**).

Tuba-seq^Ultra^ analysis indicated that *KT;H11^LSL-Cas12a^*and *KT* mice developed similar numbers of clonal expansions/tumors, consistent with the importance of oncogenic KRAS in tumorigenesis (**Figure 5g**). However, *KT;H11^LSL-Cas12a^* mice had much larger tumors and thus higher tumor burden (**Figure 5h** and **Supplementary Figure 3c-d**). Analysis of bulk tissue uncovered indels at all target sites (**Supplementary Figure 3e**). Tumors with each of the 9 crRNAs had similar tumor number and size, further underscoring the broad efficiency of Cas12a-mediated gene inactivation (**Figure 5i** and **Supplementary Figure 3f**).

Lentiviral vectors generated as a pool suffer from lentiviral template switching during reverse transcription, which can shuffle different elements^37^. We observed minimal recombination-mediated shuffling of guides between different crRNA arrays (**Supplementary Figure 4a-f**), likely due to the short distance between guides in Cas12a-based crRNA arrays, in contrast to Cas9-based approaches^37,38^. These data highlight the versatility of somatic Cas12a-mediated gene inactivation to generate novel and clinically relevant complex cancer genotypes.

## DISCUSSION

In this study, we used Cas12a mice and 3-guide crRNA arrays to rapidly generate complex genotypes in somatic cells. By synthesizing and cloning arrays in a simple pooled manner, this approach should enable the rapid generation of numerous complex genotypes in parallel, dramatically increasing both the scale and the rate at which new disease models with complex genotypes can be studied. Previous *in vitro* studies have employed as many as 25 guides together^7^, suggesting that many loci could be targeted together *in vivo* with this transgenic model.

We applied somatic CRISPR/Cas12a genome editing initially to study cancer, as cancer has tremendous genomic complexity, yet the genetically engineered mouse models that have been used to study these diseases generally have only a few designed genetic changes. While this has enabled a reductionist approach to uncover key phenotypes controlled by different genes in cancer, it has also limited our ability to model and understand key aspects of cancer that are driven by combinations of genomic alterations. For instance, engineered mouse models often fail to develop spontaneous metastases and those that do rarely metastasize to all organs commonly observed in humans^2^. Indeed, metastases tend to have a greater degree of genomic complexity than primary tumors^39^, so modeling metastatic cancer in mice may be facilitated by the generation of tumors with additional engineered alterations.

While Cre/lox- and CRISPR/Cas9-mediated somatic genome editing have been used to generate cancer models driven by complex combinations of loss of function alleles, the generation of new models remains time-consuming^2^. Furthermore, given the complexity of genotypes within human tumors, the generation of a single model of a given cancer type (*e.g.* our model of oncogene-negative lung adenocarcinoma)^20^ should not be seen as a model for all patients with that broad class of tumors. Here, we have developed enAsCas12a mice to close the gap between the complexity of human tumors and the relative simplicity of genetically engineered mouse models.

While our study focused on the use of Cas12a for multiple gene inactivation in cancer models, this system is versatile and should have numerous applications within and beyond cancer modeling. Cas12a-generated staggered DNA breaks could make it more suitable for the introduction of exogenous DNA sequences through homology-directed repair^9^. Furthermore, while we used delivery of lentiviral Cre driven by a ubiquitous promoter to initiate Cas12a expression and editing, use of Cre driven by more specific promoters would enable control of Cas12a expression in particular cell types of interest. Somatic Cas12a-mediated genome editing may also be well-suited for generating chromosomal rearrangements, such as oncogenic translocations that require the simultaneous targeting of pairs of loci, as well as the generation of panels of larger deletions. The multiplicity of Cas12a targeting could make it suitable for modeling even more complex mutational states, such as chromothripsis or aneuploidy (through chromosomal destruction). Finally, transgenic Cas12a in our models obviates the need for exogenous delivery of the large (4 kb) Cas12a gene and negates concerns of pre-existing immunity, which has been shown to dramatically affect cells with exogenous Cas9 expression^40^.

Compared to Cas9, Cas12a creates more diverse indels and continues cutting its target loci for longer due to the greater distance between its PAM and cleavage sites^41^. As a result, we previously used Cas12a to generate an evolving barcode system to map cellular phylogenies^41^. Incorporation of this system into *in vivo* models of development and disease could complement Cas9-based approaches, and the multiplexed ability of Cas12a crRNA expression could allow lineage tracing in combination with gene inactivation. This could dramatically increase our ability to both generate and study complex genotypes^42–45^.

The generation of somatic cells with complex genotypes could be of broad value, both in cancer modeling and beyond. Somatic genome editing with Cas9 transgenic mice has enabled rapid analysis of the *in vivo* effects of genetic alterations in neuroscience^46–49^, cancer^22,50–53^, immunology^54^, and other fields^55,56^. These Cas12a transgenic mice will enable the generation of pairwise and higher-order combinations of genetic alterations to be generated in cell types of interest to map complex phenotypes to complex genotypes and improve our understanding of human disease.

## METHODS

### Generation of the *H11^LSL-Cas12a^* transgenic allele

A targeting vector containing 5’ and 3’ homology arms flanking the chicken beta-actin /CMV enhancer/gamma globulin splice acceptor (CAGGS) promoter, *loxP*-STOP(6xSV40 polyA)-*loxP* cassette (LSL), enhanced *Acidaminococcus sp.* Cas12a (E174R/N282A/S542R/K548R; enAsCas12a)^13^, a nucleoplasmin nuclear localization signal (NLS), a 3xHA epitope sequence, an SV40 NLS^19^, and a rabbit β-globin polyadenylation signal (RBG pA) was used to generate the *H11^LSL-Cas12a^* knock-in mice. An sgRNA targeting the mouse *Hipp11* (*H11*) locus (GAACACTAGTGCACTTATCCTGG), the targeting vector, and Cas9 mRNA were co-injected into fertilized C57BL/6J mouse oocytes (Cyagen Biosciences). F0 founder animals were identified by PCR followed by sequence analysis. A founder mouse was bred with C57BL/6J mice to establish the *H11^LSL-Cas12a^* line. *H11^LSL-Cas12a^* mice were crossed to CMV-Cre “deleter” mice to generate the constitutive *H11^Cas12a^* mice. PCR genotyping for the *H11^LSL-Cas12a^* transgene was performed with the following primers: forward, 5’ ATGCCATCATGCTCTCACTGC 3’; reverse, 5’ GGCTATGAACTAATGACCCCGTAATTG 3’; alternative reverse, 5’ CTTGTGGGTCTTCCACCTTTCTT 3’. PCR genotyping to distinguish *H11^LSL-Cas12a^*from *H11^Cas12a^* was performed with the following primers: forward, 3’ AGGTCGAGGGACCTAATAACTTCG 5’; alternative forward, 5’ ATCTGTGCGGAGCCGAAATC 3’; reverse, 5’ TGCGTGCTTTGTCTTCCTCG 3’.

### Design, generation, barcoding, and production of lentiviral vectors

Pre-crRNA arrays with three guides were designed with diverse direct repeat sequences flanking each guide^11^, with BsmBI cut sites on both ends of the array (all guide and array sequences in **Supplementary Table 1**). Guides for enAsCas12a were designed using CRISPick for the Mouse GRCm38 reference genome (NCBI RefSeq v.108.20200622)^11,57^. Arrays were ordered as single-stranded DNA pools (Twist Biosciences) containing all 27 combinations of 3 guides for each of the 3 genes targeted in an array. Barcoded lentiviral vector backbones were created by cloning a 98 bp oligo with a 16-nucleotide diverse barcode (BC) and two BsmBI restriction sites into the 3’ end of the bovine U6 promoter with Gibson Assembly (NEBuilder HiFi, New England Biosciences) in a vector also containing PGK-Cre recombinase^23^. After low-cycle PCR amplification, pre-crRNA arrays were ligated via Golden Gate Assembly with BsmBI into the barcoded vector backbone. The cloning product was then electroporated into competent *E. coli* (C3020K, New England Biosciences) and plated onto LB-ampicillin plates. To ensure sufficient barcode diversity for Tuba-seq^Ultra^ sequencing and delineation of tumors, ∼10^6^ colonies were collected for each Lenti-U6^BC^-crRNA-Cre pool.

Lentiviral vectors were produced using polyethylenimine (PEI)-based transfection of 293T cells with delta8.2 and VSV-G packaging plasmids in 15 cm cell culture plates. Sodium butyrate (B5887, Sigma Aldrich) was added 8 hours after transfection to achieve a final concentration of 20 mM. Media was refreshed 24 hours after transfection. 20 mL of virus-containing supernatant was collected 36, 48, and 60 hours after transfection. The three collections were then pooled and concentrated by ultracentrifugation (112,000 g for 1.5 hours), resuspended overnight in 120 µL PBS, and then frozen at -80°C. Viruses were titered against a lab standard of known titer.

### Mice and tumor initiation

The use of mice for this study was approved by the Institutional Animal Care and Use Committee at Stanford University, protocol number 26696. CMV-Cre (RRID:IMSR_JAX:006054)^58^, *Kras^LSL-G12D/+^* (RRID:IMSR_JAX:008179)^59^ and *R26^LSL-tdTomato^* (RRID:IMSR_JAX:007914)^60^ mice were on a C57BL/6J background. *H11^LSL-Cas12a^*mice (JAX:038388) and *H11^Cas12a^* mice (JAX:038389) are available through the Jackson Laboratory.

Lung tumors were initiated by intratracheal delivery of 60 μL of lentiviral vectors in PBS to isoflurane-anesthetized mice. Pancreatic tumors were initiated by retrograde ductal delivery of 150 μL of lentiviral vectors in PBS to isoflurane-anesthetized mice as previous described, and mice were injected intraperitoneally with 100 µg/kg cerulein (C9026, Sigma-Aldrich) every hour for 8 hours over two consecutive days, two weeks after lentiviral delivery^36^. For small-cell lung cancer tumors, mice were pre-treated with naphthalene^53^. Briefly, corn oil (C8267, Sigma-Aldrich) was filter-sterilized with a 0.22 µm filter and aliquoted into 5 mL portions. Naphthalene was then dissolved into the corn oil vehicle at a concentration of 50 mg/mL before it was administered into mice via intraperitoneal injections at a dosage of 200 mg/kg. Roughly 46-48 hours later, mice were administered virus intratracheally and supplemented with a bowl of DietGel76A (72-07-5022, ClearH2O) food to promote recovery. Infectious units (ifu) of lentivirus used for each experiment are indicated in each respective figure.

### Histology and immunohistochemistry

Tissues were fixed in 4% formalin for 24 hours, stored in 70% ethanol, and paraffin-embedded. 4 µm thick sections were used for Hematoxylin and Eosin staining and immunohistochemistry. Immunohistochemistry was performed using an Avidin/Biotin Blocking Kit (SP-2001, Vector Laboratories), Avidin-Biotin Complex kit (PK-4001, Vector Laboratories), and DAB Peroxidase Substrate Kit (SK-4100, Vector Laboratories) following standard protocols. Alcian Blue and Trichrome stains were performed following standard protocols (Histo-Tec Laboratory, Inc.). The following primary antibodies were used: anti-HA (3724, Cell Signaling Technology), anti-TTF1 (NKX2-1; ab76013, Abcam), anti-CK19 (Ab2133570, TROMA-III, Developmental Study Hybridoma Bank), and anti-UCHL1 (HPA005993, Sigma-Aldrich).

### Western blot analyses

For immunoblotting, cells and homogenized tissues were lysed in Cell Lysis Buffer (Cell Signaling Technology) containing a complete mini-protease inhibitor cocktail (Roche), then equal quantities of protein lysate (3-10 µg) were separated by SDS-PAGE using 4–12% gradient gels (Invitrogen) and transferred to PVDF membranes. Membranes were blocked with 5% milk in PBS with 0.1% Tween 20 (PBST) for 1 hour at room temperature, followed by incubation with primary antibodies diluted in PBST with 5% milk overnight at 4°C. After 4x15 minute washes with PBST, membranes were incubated with an HRP-conjugated goat anti-rabbit secondary antibody (12-348, Sigma-Aldrich) diluted 1:5000 in PBST with 5% milk. After 4x15 minute washes with PBST, protein expression was then visualized with enhanced chemiluminescence reagents (PI80196, Fisher Scientific). Primary antibodies were used at the following dilutions: rabbit anti-HA, 1:1000 (3724, Cell Signaling); rabbit anti-Cas12a, 1:1000 (19984, Cell Signaling); and rabbit anti-alpha-Tubulin, 1:2000 (2144, Cell Signaling).

### Analysis of indels at target sites

The target sites for each crRNA were PCR-amplified from genomic DNA extracted from bulk tumor-bearing tissue (**Figure 2h** and **Supplementary Figure 3e**) or microdissected tumors (**Figure 4h**) using GoTaq Green® Mastermix (Promega) and the primer pairs listed in **Supplementary Table 2**. Microdissected tumors were isolated under a dissecting microscope; however, due to the high tumor burden, samples may have contained more than one tumor.

Amplicons were run on 1% agarose gels, gel-extracted (QIAquick PCR & Gel Cleanup Kit, Qiagen), and Sanger sequenced (Elim Biopharm, Inc). The Sanger sequencing traces were analyzed with TIDE^61^ to estimate the percent of the DNA with indels at each site. As these lungs each contain many clonal tumors, the DNA should be a mix of that from non-neoplastic cells (normal lung and stromal cells), tumors with indels at that site, and tumor with indels at the target sites of the other crRNAs targeting that gene.

### Tuba-seq^Ultra^ library preparation and analysis

Benchmark control cell lines containing unique barcodes (50,000 cells each) were added to each lung sample prior to lysis to enable the calculation of the absolute number of neoplastic cells in each tumor^23^. Bulk tumor-bearing tissues were homogenized in 6 mL of cell lysis buffer (Puregene, 158063, Qiagen) using a FastPrep-24 5G tissue homogenizer (116005500, MP Biomedicals) and digested overnight with proteinase K (AM2544, Life Technologies). To remove RNA, samples were incubated with RNase A prior to genomic DNA extraction using a Puregene kit according to the manufacturer’s recommended protocol. Q5 High-Fidelity 2x Master Mix (M0494X, New England Biolabs) was used to amplify the U6-BC-crRNA region from 32 μg of genomic DNA in a total reaction volume of 800 μL per sample using unique dual-indexed primers as described^23^. The concentration of the amplified barcode product in each PCR was measured using D1000 screentape reagents (5067-5582, Agilent Technologies) and the Tapestation instrument (Agilent Technologies). Amplicons were pooled at equal molar ratios of barcode product, normalized to the estimated burden of tumors in each mouse lung sample (measured lung mass minus an estimated normal lung weight of 0.15 g). Pooled PCR products were purified using Agencourt AMPure XP beads (A63881, Beckman Coulter) using a double size selection protocol and sequenced on the Illumina NextSeq 6000 platform (read length 2x150bp) for barcode analysis and MiSeq (2x300) for crRNA array analysis.

Paired-end reads were processed with regular expressions to identify the three gRNA sequences (henceforth referred to as the tumor-genotype) and clonal barcodes. When identifying gRNA sequences, we strictly required a perfect match with the designed sequences. The 12-nucleotide random clonal barcode sequence possesses a high theoretical diversity of approximately 4^12^ (> 10^7^). Given the virus titer we used (2x10^5^ or 1x10^6^ infectious units per mouse), there are typically fewer than 10,000 clonal tumors for each tumor-genotype per mouse. As such, the probability of two clonal tumors with the same tumor-genotype possessing two barcodes within a 1-hamming distance of each other is extremely low. Consequently, when we encountered low-frequency clonal barcodes within a 1-hamming distance of high-frequency clonal barcodes, we attributed them to sequencing or PCR errors. These low-frequency barcodes were merged with barcodes of higher frequencies. After extracting barcode and crRNA information, we converted the read counts associated with clonal tumors into absolute neoplastic cell numbers. This conversion was accomplished by normalizing the reads of the clonal tumor to the number of reads of the “spike-in” benchmark cell lines added to each sample prior to lung lysis and DNA extraction. We imposed a minimum tumor size cutoff of 300 cells for downstream analysis.

### Characterization of overall tumor growth and initiation

To quantify tumor growth, we used the log-normal mean (LN mean) tumor size, which is the maximum-likelihood estimate of mean tumor size, assuming a log-normal tumor size distribution. In addition to the tumor size metric, we characterized the effects of gene inactivation on tumorigenesis by accounting for both the number of tumors (“tumor number”) and the cumulative number of neoplastic cells (“tumor burden”). When comparing tumor growth and initiation across different genotypes of mice tumor number and tumor burden are normalized by the amount of virus delivered.

### Estimation of tumor growth and initiation effects of individual crRNA arrays

To estimate the relative effect of individual crRNAs on tumor growth, we calculated the relative LN mean tumor size by normalizing the LN mean of tumors with a specific crRNA to the average LN mean size of the three crRNAs targeting the same gene. This normalization process extended to the analysis of tumors with specific crRNA arrays, where the relative LN mean tumor size for a tumor with an array was normalized to the mean of all 27 distinct crRNA arrays. Unlike the LN mean tumor size, tumor number is linearly affected by lentiviral titer and is thus sensitive to underlying differences in the representation of each Lenti-U6^BC^-crRNA-Cre vector in the viral pool. Relative tumor number was computed by initially standardizing the tumor number in Cas12a-expressive mice against that in control mice lacking Cas12a. Subsequently, this standardized tumor number was then normalized using the same methodology applied to the relative LN mean.

### Tumor exclusion based on PCR template shuffling and unexpected crRNA arrays

Ideally, each clonal barcode would be uniquely paired with one crRNA array. However, instances where the same clonal barcode is associated with multiple crRNA arrays have been observed, which may be attributed to several factors: (1) multiple copies of the same barcode molecule ligated to distinct arrays during vector cloning; (2) template switching during lentiviral reverse transcription; and (3) PCR uncoupling throughout library preparation^37,38^. The first two scenarios would yield tumors that accurately reflect the genotype information conveyed by the crRNA array, while the latter scenario would generate spurious tumors (i.e. BC-crRNA reads that do not originate from a genuine tumor in the sample). Such spurious tumors, resulting from PCR uncoupling (a relatively rare event) will typically be markedly smaller than their genuine counterparts. To discern spurious tumors from genuine tumors with the same clonal barcode, we created a reference null distribution of size ratios between large and small tumors by randomly sampling pairs of tumors within the same mouse. When a clonal barcode was found associated with multiple crRNA arrays, we designated the crRNA array with the largest tumor as genuine and others as potentially aberrant. By calculating the size ratio of the genuine to the potential spurious tumor and comparing it against the upper 95^th^ percentile of the reference distribution, we can identify and eliminate most, if not all, spurious tumors.

In cases when tumors were found to be associated with crRNA arrays that were not from the “correct” lentiviral libraries, these tumors could have arisen from the coincident transduction of a cell with the “correct” virus and a “contaminating” virus. To eliminate these tumors from the analysis in the most conservative manner, we identified each clonal barcode linked to the “contaminating” vector, a corresponding clonal barcode associated with the “correct” vector that exhibited the most similar tumor size within the same mouse, and excluded both clonal barcodes from further analysis.

### Cell line generation

Tail tip fibroblasts were generated by mechanically dissociating depilated tail tips, digesting them in 0.25% trypsin (Life Technologies) for 30 minutes, collecting tissue fragments with centrifugation at 500 rcf for 5 minutes, then resuspending and plating in high glucose Dulbecco’s Modified Eagle Medium (Life Technologies) containing 20% fetal bovine serum and 2% penicillin-streptomycin (Thermo Fisher Scientific).

## ACKNOWLEDGEMENTS

We thank the Stanford Veterinary Animal Care Staff for expert animal care, Greg Charville for advice on histologic characterization of tumors, and members of the Winslow laboratory for helpful comments. J.D.H was supported by an American Cancer Society Fellowship (PF-21-112-01-MM) and a TRDRP Postdoctoral fellowship (T31FT1619). Y.J.T was partly supported by the Canadian Institute of Health Research (CIHR) postdoctoral fellowship (MFE-176568). P.A.R. was supported by the National Science Foundation Graduate Research Fellowship (DGE-2146755) and the Lucille P. Markey Stanford Graduate Fellowship. This work was supported by NIH R01-CA230025 (to M.M.W), NIH P01-CA244114 (to M.M.W.), NIH R01-CA231253 (to M.M.W and D.A.P), NIH R01-CA234349 (to M.M.W and D.A.P.), NIH R35-CA231997 (to J.S.), NIH R35-HG011316 and R01-GM141627 (to L.C.), and in part by the Stanford Cancer Institute support grant (NIH P30-CA124435).

---

**Supplementary Figure 1.**
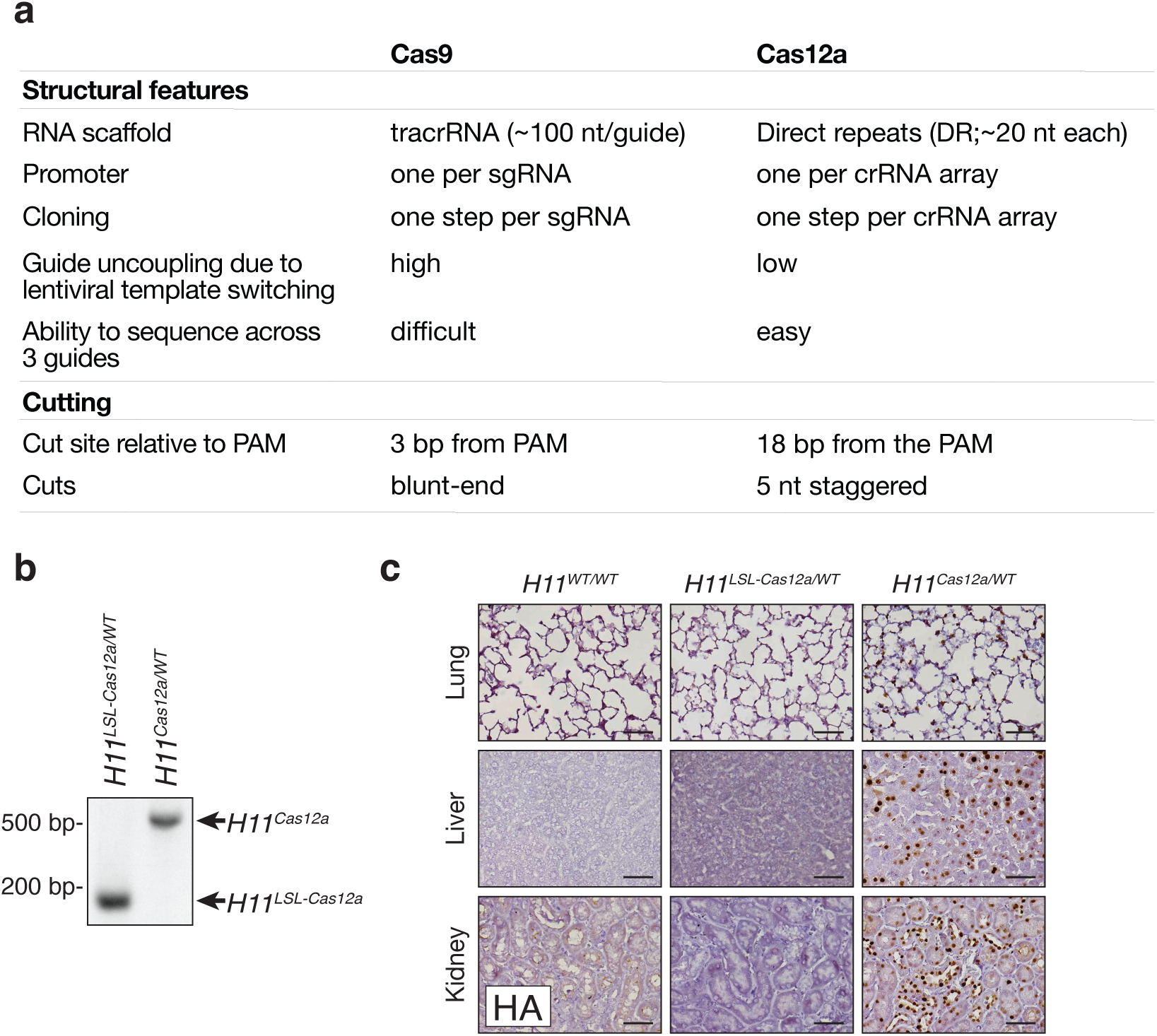
Comparison of salient features of Cas9 and Cas12a, and broad nuclear expression of Cas12a in *H11^Cas12a^* mice. **a.** Summary of features of Cas9 and Cas12a relevant to multiplexed genome editing. Note that the potential impact of off-target effects is increased when targeting more genes. **b.** PCR genotyping of mice of the indicated *H11^LSL-Cas12a^* and *H11^Cas12a^* genotypes. **c.** Immunohistochemical staining for the Cas12a HA tag in lung, liver and kidney sections from the indicated genotypes of mice. Scale bars, 50 µm. Higher magnification images of these liver sections is shown in Figure 1f.

**Supplementary Figure 2.**
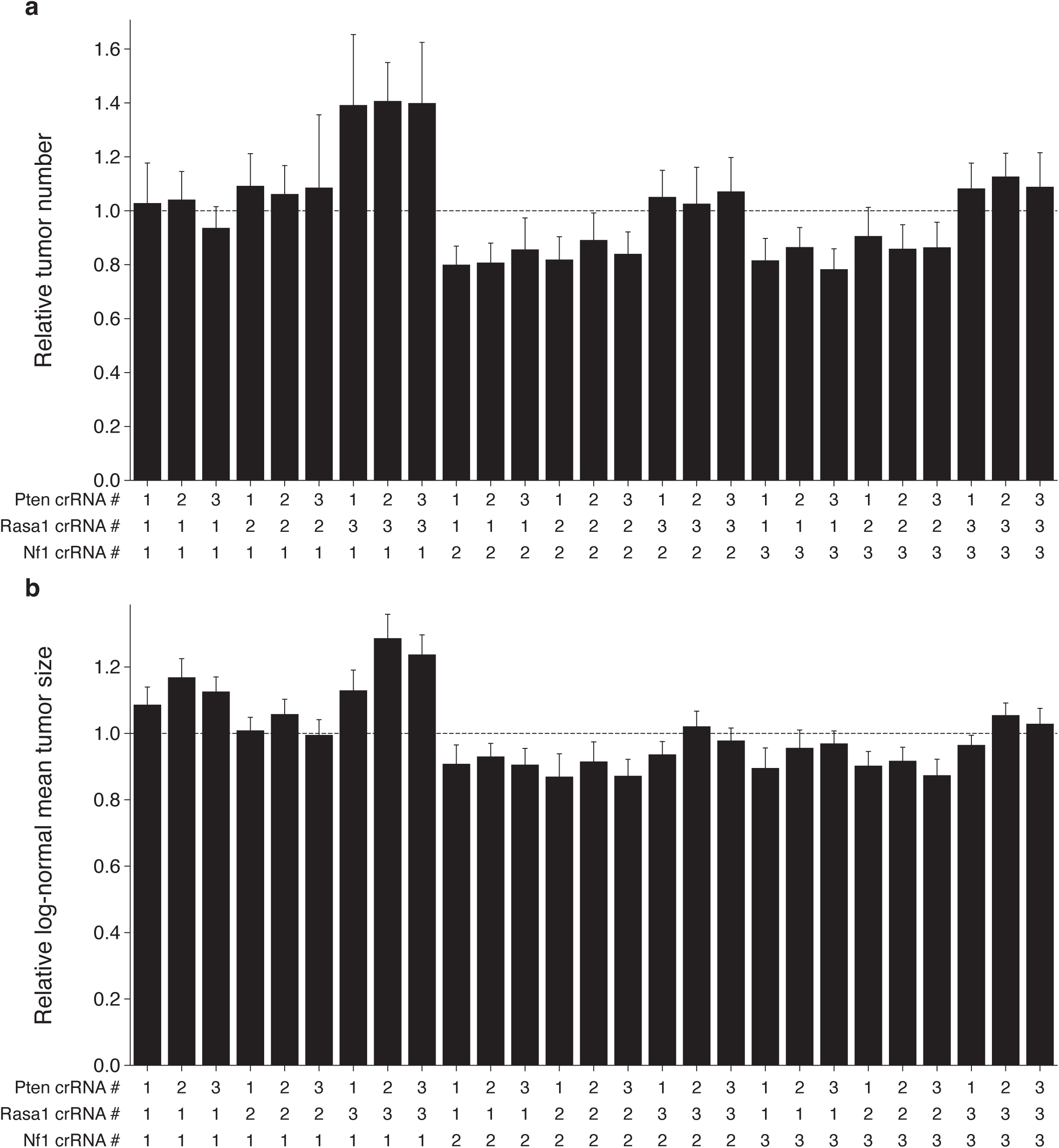
Incorporation of Cas12a-mediated genome editing with Tuba-seq^Ultra^ enables quanitification of the effects of each crRNA array. **a,b.** Number of tumors (**a**) and mean tumor size assuming a log-normal distribution (**b**) for each crRNA array (27 combinations in total) relative to the median value for all arrays in *H11^LSL-Cas12a^* mice. 95% confidence intervals are shown.

**Supplementary Figure 3.**
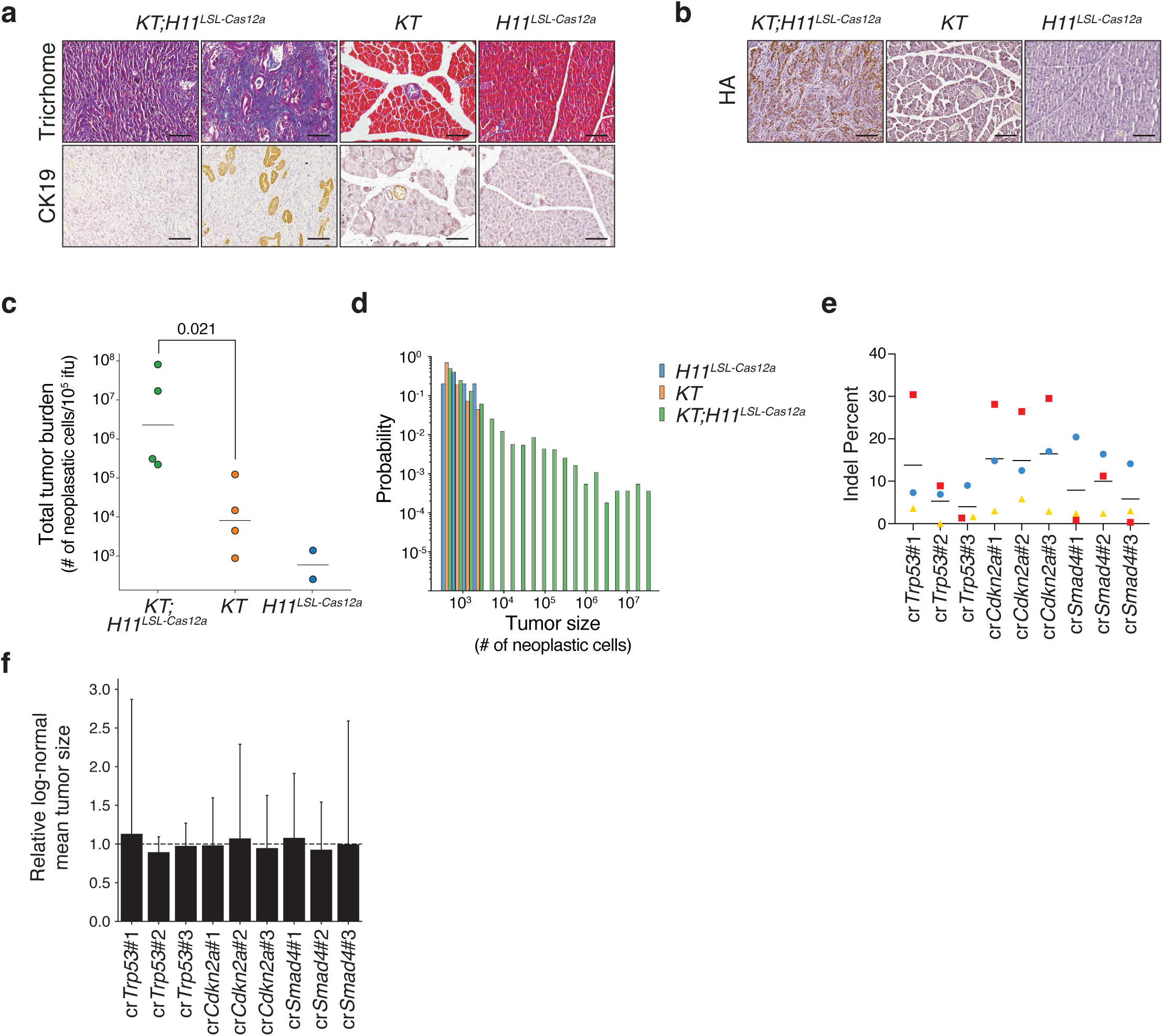
Inactivation of *Trp53*, *Cdkn2a*, and *Smad4* greatly increases tumor size and leads to the development of PDAC with differentiated areas with CK19+ cancer cells and desmoplastic stroma as well as more poorly differentiated areas. a. Immunohistochemistry images of pancreas sections from the indicated genotypes, showing Trichrome (upper) and Cytokeratin-19 (CK19; lower) staining. *KT;H11^LSL-Cas12a^* panels show representative poorly differentiated (left) and more differentiated (right) regions. Scale bars, 100 μm. b. Immunohistochemical staining for the Cas12a HA tag on pancreas tissue from representative *KT;H11^LSL-Cas12a^*, *KT* and *H11^LSL-Cas12a^* mice transduced with Lenti-U6^BC^-cr*Trp53/Cdkn2a/Smad4*-Cre. Scale bars, 100 μm. c. Total number of neoplastic cells (Total tumor burden) in each mouse normalized to viral titer. Each dot represents a mouse and the bar is the mean. Note that two *H11^LSL-Cas12a^* mice did not have detectable tumor burden and are not plotted. P-value calculated with Wilcoxon rank-sum test. d. Probability distribution of the size of tumors in the indicated genotypes of mice. e. Indel frequencies for genomic regions targeted by each crRNA within bulk tumor-bearing pancreas tissue from KT;*H11^LSL-Cas12a^*mice. Tissue samples had variable tumor burden, likely explaining observed variability in indel frequencies. Symbol shapes and colors identify values from the same mouse. Bars represent means. f. Mean tumor size assuming a log-normal distribution for each crRNA (guide) relative to the median value for each gene. 95% confidence intervals are shown.

**Supplementary Figure 4.**
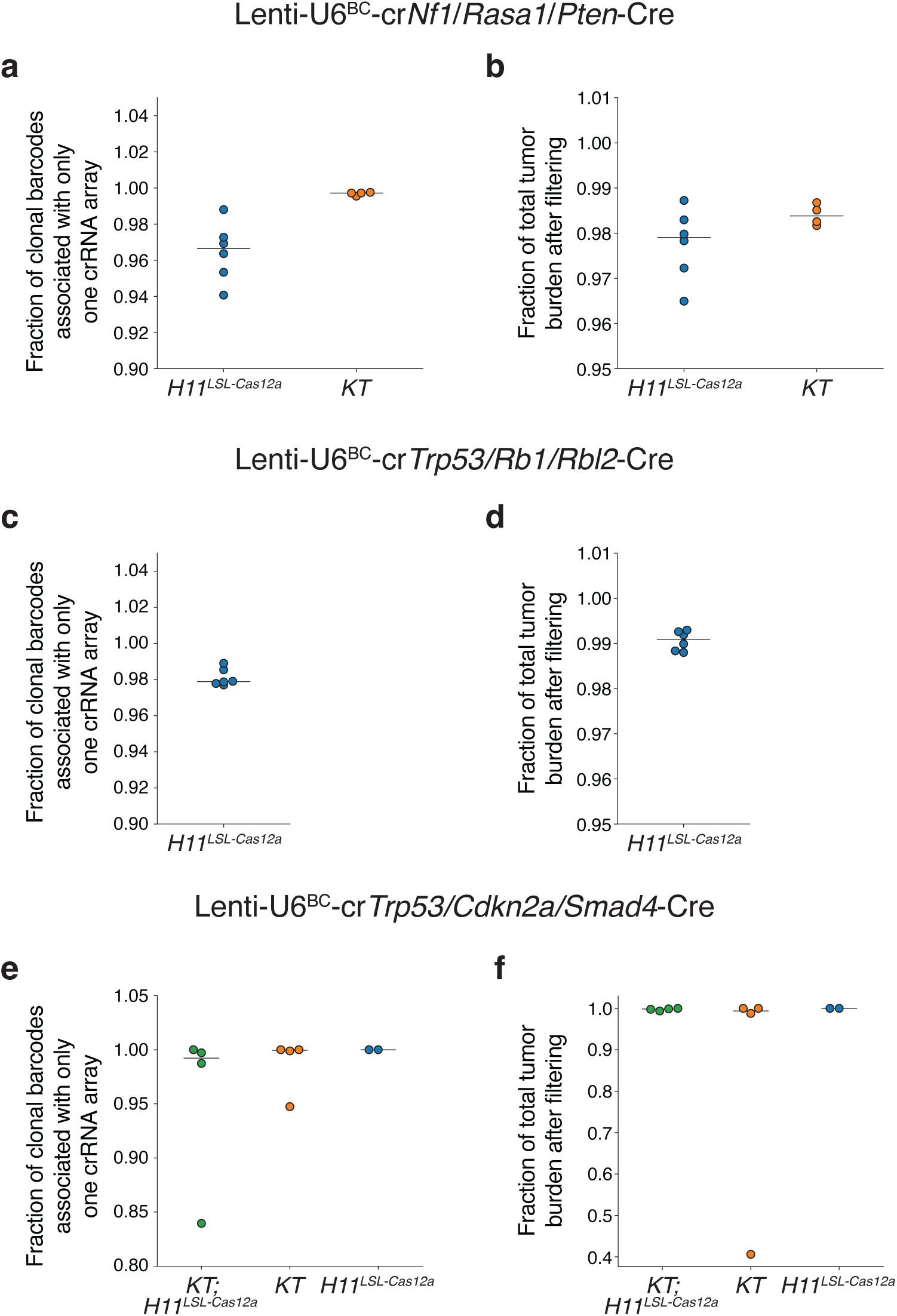
Generation of Cas12a Lenti-U6^BC^-crRNA-Cre vector libraries is associated with minimal shuffling of crRNA sequences. **a,c,e.** Fraction of all U6-integrated barcode reads that are associated with only a single crRNA array within each vector pool in the indicated mouse genotypes: Lenti-U6^BC^-cr*Nf1*/*Rasa1*/*Pten*-Cre (**a**), Lenti-U6^BC^-cr*Trp53/Rb1/Rbl2*-Cre (**c**), and Lenti-U6^BC^-cr*Trp53/Cdkn2a/Smad4*-Cre (**e**). Bars represent medians. **b,d,f.** Fraction of total neoplastic cells (total tumor burden) remaining after filtering out spurious “tumor” reads arising during library preparation and sequencing of each pool in the indicated mouse genotypes: Lenti-U6^BC^-cr*Nf1*/*Rasa1*/*Pten*-Cre (**b**), Lenti-U6^BC^-cr*Trp53/Rb1/Rbl2*-Cre (**d**), and Lenti-U6^BC^-cr*Trp53/Cdkn2a/Smad4*-Cre (**f**). Bars represent medians.

